# Genetic network regulating visual acuity makes limited contribution to visually guided eye emmetropization

**DOI:** 10.1101/2021.05.04.442661

**Authors:** Tatiana V. Tkatchenko, Andrei V. Tkatchenko

## Abstract

During postnatal development, the eye undergoes a refinement process whereby optical defocus guides eye growth towards sharp vision in a process of emmetropization. Optical defocus activates a signaling cascade originating in the retina and propagating across the back of the eye to the sclera. Several observations suggest that visual acuity might be important for optical defocus detection and processing in the retina; however, direct experimental evidence supporting or refuting the role of visual acuity in refractive eye development is lacking. Here, we used genome-wide transcriptomics to determine the relative contribution of the retinal genetic network regulating visual acuity to the signaling cascade underlying visually guided eye emmetropization.

Our results provide evidence that visual acuity is regulated at the level of molecular signaling in the retina by an extensive genetic network. The genetic network regulating visual acuity makes relatively small contribution to the signaling cascade underlying refractive eye development. This genetic network primarily affects baseline refractive eye development and this influence is primarily facilitated by the biological processes related to melatonin signaling, nitric oxide signaling, phototransduction, synaptic transmission, and dopamine signaling. We also observed that the visual-acuity-related genes associated with the development of human myopia are chiefly involved in light perception and phototransduction. Our results suggest that the visual-acuity-related genetic network primarily contributes to the signaling underlying baseline refractive eye development, whereas its impact on visually guided eye emmetropization is modest.

## 1. Introduction

The general layout of the eye is established during embryonic development; however, the eye undergoes a refinement process during postnatal period, whereby the optical geometry of the eye and its refractive state are guided towards sharp vision in a process called emmetropization [1–8]. Various environmental factors influence refractive eye development during emmetropization [8–11]; however, the leading environmental factor guiding refractive eye development is optical defocus [8, 12]. The eye is very sensitive to the sign of optical defocus and can accurately compensate for imposed defocus by modulating the growth of the posterior segment of the eye [12–18]. Emmetropization uses optical defocus to match the eye’s axial length to its focal plane and normally produces emmetropia (sharp vision) [8]. Prolonged exposure to positive optical defocus leads to the development of hyperopia (farsightedness), whereas excessive exposure to negative optical defocus leads to the development of myopia (nearsightedness) [13, 14, 16–27]. The role of optical defocus in eye emmetropization is well established [8]. It was also found that molecular signaling underlying retinal response to positive and negative defocus propagate along two largely distinct signaling cascades [12]. Genetic networks underlying baseline refractive eye development and visually guided eye emmetropization are also beginning to come to light [8, 28, 29]. However, the molecular cascade responsible for the detection and processing of optical defocus by the eye is still poorly understood. We have recently found that the gene expression network subserving perception of contrast plays an important role in optical defocus detection and visually guided eye emmetropization [30]; however, the role of visual acuity in optical defocus detection and emmetropization remains a subject of debate.

Visual acuity is primarily determined by the density of photoreceptors in the central retina and declines sharply with increasing retinal eccentricity [31, 32]. Several lines of evidence suggest that the absolute level of visual acuity does not play a significant role in optical defocus detection and emmetropization because species with different levels of visual acuity ranging from 1.3 cycles per degree (cpd) in fish [33], 0.6-1.4 cpd in mice [34–36], 2.4 cpd in tree shrews [37, 38], 2.7 cpd in guinea pigs [39, 40], 5 cpd in cats [41, 42], 5 cpd chickens [43, 44] and 44 cpd in rhesus monkeys and humans [45–47] undergo emmetropization and can compensate for imposed optical defocus [13–17, 48–52]. Interestingly, visual acuity in newborn human infants during first months of life, i.e., when they undergo most active eye emmetropization, remains low, 2.4 cpd in 1-month-old infants gradually increasing to 10-15 cpd in 8-month-olds [47, 53, 54]. In agreement with these observations, Mutti et al. found that emmetropization in infants does not appear to be dependent on visual acuity [55]. Studies in a mutant chicken model with diminished visual acuity also revealed that reduced visual acuity had limited impact on susceptibility to either form-deprivation or lens-induced myopia, hence emmetropization [56]. It was also found that accommodation and emmetropization are primarily dependent on low spatial frequencies even in species with high visual acuity [57, 58]; and that the peripheral retina, which has much lower visual acuity than the central retina [32, 53, 59–62], plays an important role in defocus detection and emmetropization [8, 63–65]. Moreover, it was reported that inhibitory inputs to the ganglion cells from the amacrine cells, which are critical for optical defocus detection, are more predominant in the low-resolution peripheral retina, while the high-resolution foveal retinal circuits primarily rely on excitatory inputs [66]. Nevertheless, there is evidence that visual acuity might influence visually guided eye emmetropization. For example, certain mutations affecting visual acuity in humans were also found to be linked to the development of myopia [67]. It was also found that maturation of visually evoked potentials (VEPs), which correlate with sensitivity to optical defocus, shows strong correlation with visual acuity [68].

Similar to refractive eye development, visual acuity is controlled by both environmental and genetic factors. Prusky et al. discovered that environmental enrichment during early postnatal period, such as exposure to stationary and moving visual patterns, enhances visual acuity, whereas visual deprivation reduces it [69, 70]. Genetic background also has a significant impact on visual acuity. For example, different inbred strains of mice raised under the same environmental conditions exhibit large differences in visual acuity and ability to discriminate patterns [71]. Genetic polymorphism in the gene encoding complement factor H (*CFH*) was found to affect visual acuity in humans [72]. It was also found that mutations in genes encoding retinal synaptic proteins, such as the *blu* gene encoding a vesicular glutamate transporter and *Lrit1* encoding a leucine-rich transmembrane protein, affect visual acuity without changing the density of photoreceptors [73, 74]. Glucose metabolism in the retina has been implicated in the regulation of visual acuity [75, 76]. Finally, cannabinoids were shown to influence visual acuity in mice via the cannabinoid receptor type 2 (CB2R) [77]. All this evidence suggests that visual acuity is controlled by a genetic network, which integrates visual experience with the downstream molecular signaling in the retina. Considering that some data suggest that visual acuity might be linked to refractive eye development, it is critical to characterize the genetic network and signaling pathways subserving visual acuity and determine their contribution to baseline refractive eye development and visually guided eye emmetropization.

Here, we analyzed gene expression networks and signaling pathways regulating visual acuity in a panel of genetically diverse mice, which have different visual acuity, different baseline refractive errors, and different susceptibility to form-deprivation myopia, and determined the contribution of the genetic network subserving visual acuity to baseline refractive eye development and visually guided eye emmetropization.

## 2. Materials and methods

### 2.1. Ethics statement

Inbred mouse strains (129S1/svlmj, A/J, C57BL/6J, CAST/EiJ, NOD/ShiLtJ, NZO/HlLtJ, PWK/PhJ, and WSB/EiJ) were obtained from the Jackson Laboratory (Bar Harbor, ME) and were maintained as an in-house breeding colony. Food and water were provided ad libitum. All procedures adhered to the Association for Research in Vision and Ophthalmology (ARVO) statement on the use of animals in ophthalmic and vision research and were approved by the Columbia University Institutional Animal Care and Use Committee. Animals were anesthetized via intraperitoneal injection of ketamine (90 mg/kg) and xylazine (10 mg/kg) and were euthanized using CO_2_ followed by cervical dislocation. The study was carried out in compliance with the ARRIVE guidelines.

### 2.2. Analysis of visual acuity

Visual acuity was measured at P40 using a virtual optomotor system (Mouse OptoMotry System, Cerebral Mechanics) (Supplementary Table S1), as previously described [78]. Briefly, the animal to be tested was placed on a platform surrounded by four computer screens displaying a virtual cylinder comprising a vertical sine wave grating in 3D coordinate space. The OptoMotry software controlled the speed of rotation, direction of rotation, the frequency of the grating and its contrast. To measure visual acuity, the initial spatial frequency of the grating was set at 0.1 cycles/degree (cpd) and the contrast was set at maximum. The frequency was then systematically increased using staircase procedure until the maximum spatial frequency capable of eliciting a response (visual acuity) was determined. The staircase procedure was such that 3 correct answers in a row advanced it to a higher spatial frequency, while 1 wrong answer returned it to a lower frequency.

### 2.3. Analysis of baseline refractive error

We measured baseline refractive errors in both left and right eyes on alert animals at P40 using an automated eccentric infrared photorefractor as previously described (Supplementary Table S2) [79, 80]. The animal to be refracted was immobilized using a restraining platform, and each eye was refracted along the optical axis in dim room light (< 1 lux), 20-30 minutes after the instillation of 1% tropicamide ophthalmic solution (Alcon Laboratories) to ensure mydriasis and cycloplegia. Refractive error measurements were automatically acquired by the photorefractor every 16 msec. Each successful measurement series (i.e., Purkinje image in the center of the pupil and stable refractive error for at least 5 sec.) was marked by a green LED flash, which was registered by the photorefractor software. Five independent measurement series were taken for each eye. Sixty individual measurements from each series, immediately preceding the green LED flash, were combined, and a total of 300 measurements (60 measurements x 5 series = 300 measurements) were collected for each eye. Data for the left and right eyes were combined (600 measurements total) to calculate the mean refractive error and standard deviation for each animal.

### 2.4. Analysis of susceptibility to form-deprivation myopia

We measured the extent of myopia induced by diffuser-imposed retinal image degradation (visual form deprivation) (Supplementary Table S3). The eye is the most sensitive to optical defocus during a critical period in postnatal development, which continues in mice from the eye opening (P12-P14) to approximately P60 [49, 79]. Therefore, visual form deprivation was induced in P24 mice by applying plastic diffusers to one of the eyes and refractive development of the treated eye was compared to that of the contralateral eye (contralateral eye was not treated with a diffuser) as previously described [49, 81]. Diffusers represented low-pass optical filters, which degraded the image projected onto the retina by removing high spatial frequency details. Hemispherical plastic diffusers were made from zero power rigid contact lenses manufactured from OP3 plastic (diameter = 7.0 mm, base curve = 7.0 mm; Lens.com) and Bangerter occlusion foils (Precision Vision). Diffusers were inserted into 3D-printed plastic frames (Proto Labs). On the first day of the experiment (P24), animals were anesthetized via intraperitoneal injection of ketamine and xylazine, and frames with diffusers were attached to the skin surrounding the eye with six stitches using size 5-0 ETHILON™ microsurgical sutures (Ethicon) and reinforced with Vetbond™ glue (3M Animal Care Products) (the contralateral eye served as a control). Toenails were covered with adhesive tape to prevent mice from removing the diffusers. Animals recovered on a warming pad and were then housed under low-intensity constant light in transparent plastic cages for the duration of the experiment as previously described [49, 81]. Following 21 days of visual form deprivation (from P24 through P45), diffusers were removed and the refractive state of both treated and control eyes was assessed using an automated infrared photorefractor as previously described [79, 80]. The interocular difference in refractive error between the treated and contralateral control eye was used as a measure of induced myopia.

### 2.5. RNA extraction and RNA-seq

Animals were euthanized following an IACUC-approved protocol. Eyes were enucleated, the retinae were dissected from the enucleated eyes. The retinae were washed in RNAlater (Thermo Fisher Scientific) for 5 min., frozen in liquid nitrogen, and stored at −80°C. To isolate RNA, tissue samples were homogenized at 4°C in a lysis buffer using Bead Ruptor 24 tissue homogenizer (Omni). Total RNA was extracted from each tissue sample using miRNAeasy mini kit (QIAGEN) following the manufacturer’s protocol. The integrity of RNA was confirmed by analyzing 260/280 nm ratios (Ratio_260/280_ = 2.11-2.13) on a Nanodrop (Thermo Scientific) and the RNA Integrity Number (RIN = 9.0-10.0) using Agilent Bioanalyzer. Illumina sequencing libraries were constructed from 1 μg of total RNA using the TruSeq Stranded Total RNA LT kit with the Ribo-Zero Gold ribosomal RNA depletion module (Illumina). Each library contained a specific index (barcode) and were pooled at equal concentrations using the randomized complete block (RCB) experimental design before sequencing on Illumina HiSeq 2500 sequencing system. The number of libraries per multiplexed sample was adjusted to ensure sequencing depth of ~70 million reads per library (paired-end, 2 x 100 bp). The actual sequencing depth was 76,773,554 ± 7,832,271 with read quality score 34.5 ± 0.4.

### 2.6. Post-sequencing RNA-seq data validation and analysis

The FASTQ raw data files generated by the Illumina sequencing system were imported into Partek Flow software package (Partek), libraries were separated based on their barcodes, adapters were trimmed and remaining sequences were subjected to pre-alignment quality control using the Partek Flow pre-alignment QA/QC module. After the assessment of various quality metrics, bases with the quality score < 34 were removed (≤ 5 bases) from each end. Sequencing reads were then mapped to the mouse reference genome Genome Reference Consortium Mouse Build 38 (GRCm38/mm10, NCBI) using the STAR aligner resulting in 95.0 ± 0.4% mapped reads per library, which covered 35.4 ± 1.0% of the genome. Aligned reads were quantified to transcriptome using the Partek E/M annotation model and the NCBI’s RefSeq annotation file to determine read counts per gene/genomic region. The generated read counts were normalized by the total read count and subjected to the analysis of variance (ANOVA) to detect genes whose expression correlates with either visual acuity, refractive error or susceptibility to form-deprivation myopia. Differentially expressed transcripts were identified using a P-value threshold of 0.05 adjusted for genome-wide statistical significance using Storey’s q-value algorithm [82]. To identify sets of genes with coordinate expression, differentially expressed transcripts were clustered using the Partek Flow hierarchical clustering module, using average linkage for the cluster distance metric and Euclidean distance metric to determine the distance between data points. Each RNA-seq sample was analyzed as a biological replicate, thus, resulting in three (3) biological replicates per strain.

### 2.7. Gene ontology analysis and identification of canonical signaling pathways

To identify biological functions (gene ontology categories), which were significantly associated with the genes whose expression correlated with visual acuity, baseline refractive errors or susceptibility to form-deprivation myopia, we used the database for annotation, visualization and integrated discovery (DAVID) version 6.8 [83] and GOplot R package version 1.0.2 [84]. DAVID uses a powerful gene-enrichment algorithm and Gene Concept database to identify biological functions (gene ontology categories) affected by differential genes. GOplot integrates gene ontology information with gene expression information and predicts the effects of gene expression changes on biological processes. DAVID uses a modified Fisher’s exact test (EASE score) with a P-value threshold of 0.05 to estimate the statistical significance of enrichment for specific gene ontology categories. The IPA Pathways Activity Analysis module (Ingenuity Pathway Analysis, QIAGEN) was used to identify canonical pathways encoded by the genes involved in baseline refractive eye development, susceptibility to myopia or contrast perception, and to predict the effects of gene expression differences in different mouse strains on the pathways. The activation z-score was employed in the Pathways Activity Analysis module to predict activation or suppression of the canonical pathways. The z-score algorithm is designed to reduce the chance that random data will generate significant predictions. The z-score provides an estimate of statistical quantity of change for each pathway found to be statistically significantly affected by the changes in gene expression. The significance values for the canonical pathways were calculated by the right-tailed Fisher’s exact test. The significance indicates the probability of association of molecules from a dataset with the canonical pathway by random chance alone. The Pathways Activity Analysis module determines if canonical pathways, including functional end-points, are activated or suppressed based on the gene expression data in a dataset. Once statistically significant canonical pathways were identified, we subjected the datasets to the Core Functional Analysis in IPA to compare the pathways and identify key similarities and differences in the canonical pathways underlying visual acuity, baseline refractive development, susceptibility to form-deprivation myopia.

### 2.8. Identification of genes linked to human myopia and other eye-related human genetic disorders

All mouse genes, which were found to be associated with visual acuity and baseline refractive errors or susceptibility to form-deprivation myopia were analyzed for their association with eye-related human genetic disorders. To identify genes linked to human myopia, we compared the genes that we found in mice with a list of genes located in the human myopia quantitative trait loci (QTLs). We first compiled a list of all single-nucleotide polymorphisms (SNPs) or markers exhibiting a statistically significant association with myopia in the human linkage or genome-wide association studies (GWAS) using the Online Mendelian Inheritance in Man (OMIM) (McKusick-Nathans Institute of Genetic Medicine, Johns Hopkins University) and NHGRI-EBI GWAS Catalog [85] databases. The LDlink’s LDmatrix tool (National Cancer Institute) was used to identify SNPs in linkage disequilibrium and identify overlapping chromosomal loci. We then used the UCSC Table Browser to extract all genes located within critical chromosomal regions identified by the human linkage studies or within 200 kb (±200 kb) of the SNPs found by GWAS. The list of genes located within human QTLs was compared with the list of genes that we found in mice using Partek Genomics Suite (Partek). To identify genes associated with eye-related human genetic disorders unrelated to myopia, we screened mouse genes that we found in this study against the Online Mendelian Inheritance in Man (OMIM) (McKusick-Nathans Institute of Genetic Medicine, Johns Hopkins University) database.

### 2.9. Statistical data analysis and data graphing

Statistical analyses of the RNA-seq data were performed using statistical modules integrated into Partek Flow (Partek), DAVID [83] and IPA (QIAGEN) software packages and described in the corresponding sections. Other statistical analyses were performed using the STATISTICA software package (StatSoft). The statistical significance of the overlaps between gene datasets was estimated using probabilities associated with the hypergeometric distribution as implemented in the Bioconductor R software package GeneOverlap and associated functions. Data graphing and visualization was performed using Partek Flow (Partek) and IPA (QIAGEN) graphing and visualization modules, as well as Prism 8 for Windows (GraphPad) and GOplot R package [84].

## 3. Results

### 3.1. Visual acuity exhibits weak but significant association with baseline refractive error and visually guided eye emmetropization in mice

In order to determine the role of visual acuity in baseline refractive eye development and visually guided eye emmetropization, we analyzed the relationship between visual acuity, baseline refractive error, and susceptibility to form-deprivation myopia (as a proxy for visually guided eye emmetropization) in eight genetically different inbred strains of mice characterized by different levels of visual acuity, baseline refractive errors, and susceptibility to form-deprivation myopia (Supplementary Tables S1-S3). Visual acuity (VA), baseline refractive error (RE), and induced myopia (myopia) were measured in 129S1/svlmj, A/J, C57BL/6J, CAST/EiJ, NOD/ShiLtJ, NZO/HlLtJ, PWK/PhJ, and WSB/EiJ mice at P40-P45 (see Methods).

We found that all three attributes (visual acuity, refractive error, and susceptibility to myopia) were inherited as quantitative traits (Figure 1A-C; Supplementary Tables S1-S3). We also found that all three attributes were strongly influenced by genetic background (ANOVA_VA_, F(7, 57)=1807.3, P < 0.00001; ANOVA_RE_, F(7, 145) = 429.8, P < 0.00001; ANOVA_myopia_, F(7, 48) = 9.8, P < 0.00001). In agreement with our recent report [30], correlation between baseline refractive error and susceptibility to myopia was weak (r = 0.2686, P = 0.0305; Supplementary Figure S1). Linear regression analyses revealed weak but statistically significant positive correlation between visual acuity and baseline refractive error (r = 0.371, P = 0.0023) (Figure 1D), as well as between visual acuity and susceptibility to myopia (r = 0.3367, P = 0.0061) (Figure 1E). Correlation coefficients for both regressions were similar suggesting essentially equal, yet relatively modest, contribution of visual acuity to both baseline refractive eye development and visually guided eye emmetropization.

**Fig. 1.**
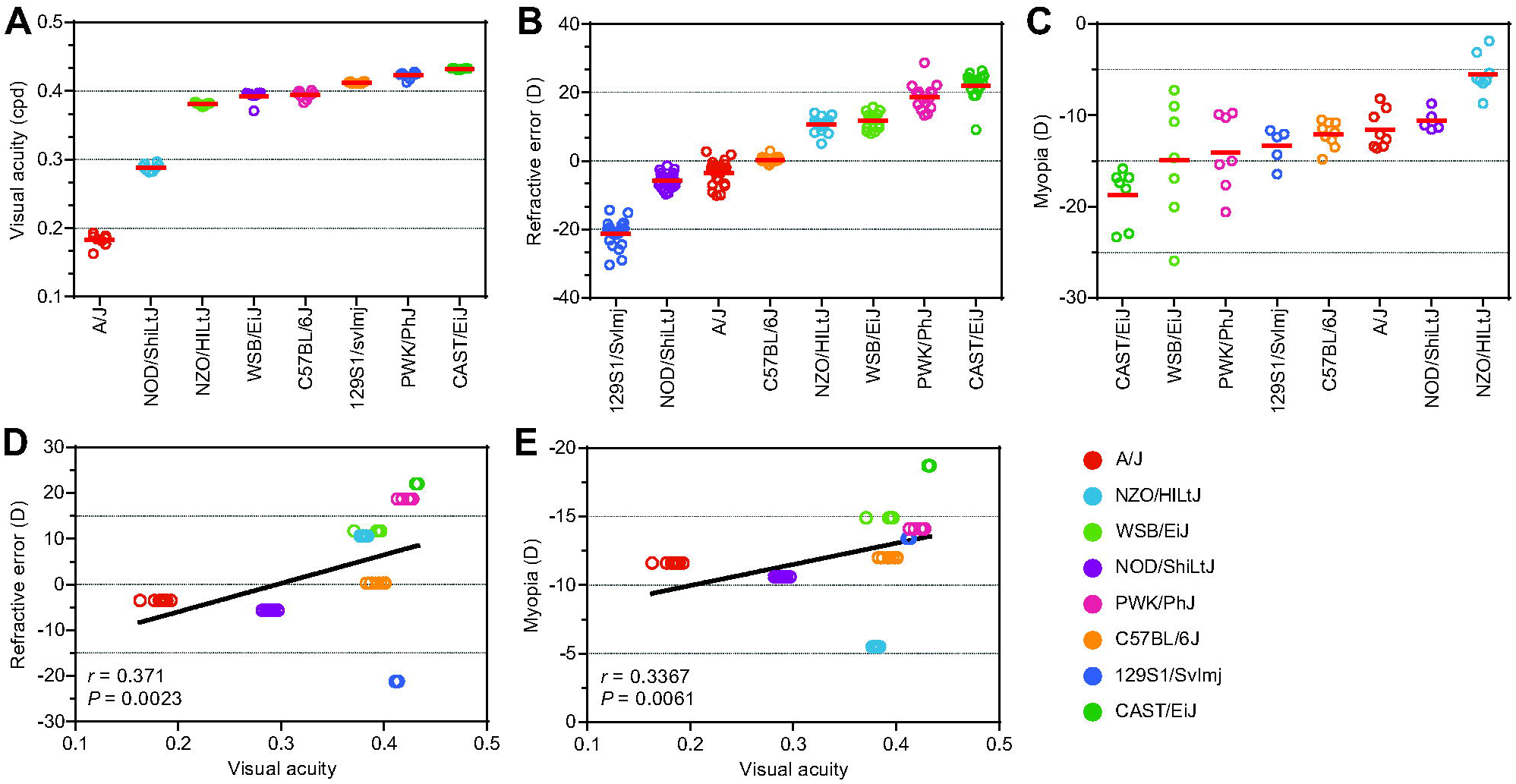
Visual acuity exhibits weak correlation with baseline refractive error and form-deprivation myopia. (A) Visual acuity is influenced by genetic background. Horizontal red lines identify means of visual acuity for each strain, while each dot represents a mean visual acuity for individual animals. (B) Baseline refractive errors range from highly myopic to highly hyperopic in mice depending on the genetic background. Horizontal red lines show mean refractive errors for each strain, while each dot corresponds to mean refractive errors of individual animals. (C) Susceptibility to form-deprivation myopia in mice is determined by genetic background. Horizontal red lines identify means of induced myopia for each strain, while each dot represents a mean myopia for individual animals. (D) Visual acuity exhibits weak statistically significant correlation with baseline refractive error. Linear regression showing weak correlation between visual acuity and baseline refractive error. r, Pearson’s correlation coefficient; P, Pearson’s correlation significance. (E) Visual acuity exhibits weak statistically significant correlation with form-deprivation myopia. Linear regression showing weak correlation between visual acuity and myopia. r, Pearson’s correlation coefficient; P, Pearson’s correlation significance.

### 3.2. Visual acuity is controlled by an extensive genetic network

Our data suggested that visual acuity is weakly associated with both baseline refractive eye development and visually guided eye emmetropization. Therefore, we then set out to investigate whether the signaling cascade underlying visual acuity is involved in the regulation of baseline refractive eye development and eye emmetropization. To gain insight into the signaling cascade subserving visual acuity and examine its involvement in the regulation of baseline refractive eye development and eye emmetropization, we performed a genome-wide gene expression profiling in the retina of the eight mouse strains listed above and analyzed the transcriptomes underlying visual acuity. The retinal acuity-related transcriptome was analyzed at P28 (age when refractive eye development in mice is progressing towards its stable plateau, but visual input is still influencing eye growth [49, 79]) using RNA-seq.

We found that visual acuity in mice correlates with expression of 1,073 retinal genes (Q-value < 0.02) (Figure 2A, Supplementary Table S4). Expression of 394 of these genes was positively correlated with visual acuity (i.e., expression was increased in animals with high visual acuity and reduced in animals with low visual acuity), whereas expression of 679 of these genes was negatively correlated with visual acuity (i.e., expression was decreased in animals with high visual acuity and increased in animals with low visual acuity). Furthermore, visual acuity was associated with biological processes related to protein phosphorylation, homophilic cell adhesion, metabolic process, intracellular signal transduction, transcriptional regulation of gene expression, axonogenesis, response to ischemia, and photoreceptor cell maintenance, among other processes (Figure 2B, Supplementary Table S7). We also found that visual acuity was linked to the activation or suppression of several canonical signaling pathways, including protein kinase A signaling, RhoGDI signaling, oleate biosynthesis, cAMP-mediated signaling, melatonin signaling, senescence pathway, Wnt/Ca+ pathway, opioid signaling pathway, prolactin signaling, PPARα/RXRα activation pathway, PTEN signaling, nNOS signaling, and β-adrenergic signaling (Figure 2C, Supplementary Table S10).

**Fig. 2.**
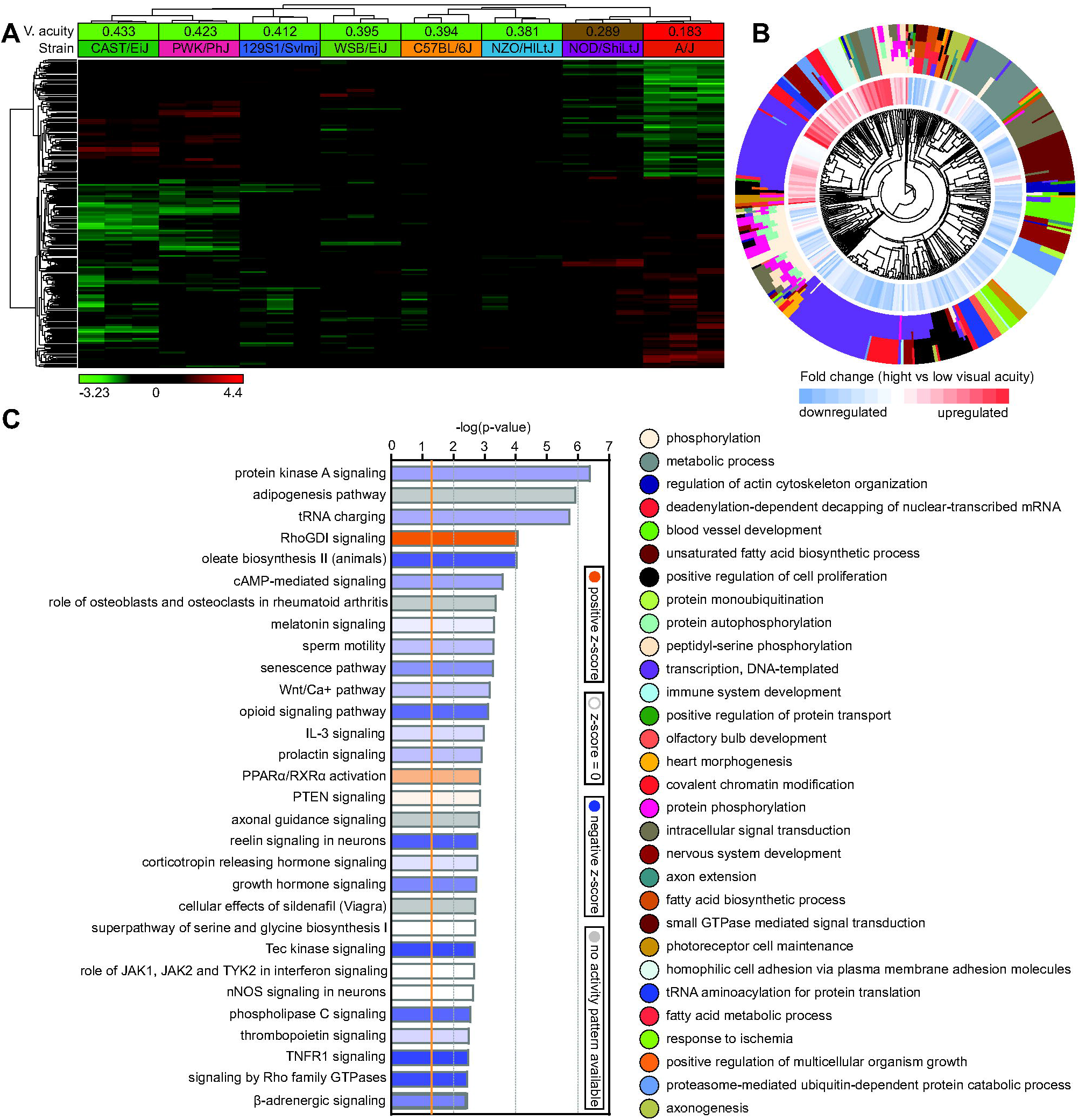
Gene expression network regulating visual acuity controls diverse biological functions and signaling pathways. (A) Expression of 1,073 genes (Q-value < 0.02) correlates with visual acuity. Hierarchical clustering shows that differential genes are organized in two clusters: one (top) cluster exhibits increased expression in mice with high visual acuity, and the second (bottom) cluster shows increased expression in mice with low visual acuity. (B) Top 30 biological processes affected by the genes involved in the regulation of visual acuity. Outer circle of the hierarchical clustering diagram shows hierarchical clusters of biological processes (identified by different colors); inner circle shows clusters of the associated genes up- or down-regulated in mice with high visual acuity versus mice with low visual acuity. (C) Top 30 canonical pathways affected by the genes involved in the regulation of visual acuity. Horizontal yellow line indicates P = 0.05. Z-scores show activation or suppression of the corresponding pathways in animals with high visual acuity versus animals with low visual acuity.

These data suggest that visual acuity is regulated by an extensive genetic network encoding a diverse set of biological processes and signaling pathways.

### 3.3. Comparison of the transcriptome for visual acuity with the transcriptome for baseline refractive eye development reveals significant contribution of the genetic network subserving visual acuity to baseline refractive development

To examine the contribution of the genetic network regulating visual acuity to baseline refractive eye development, we analyzed the overlap between gene expression networks underlying visual acuity and baseline refractive development. As reported previously [30], we identified 2,050 genes (Q-value < 0.01) whose expression was associated with baseline refractive eye development (Supplementary Figure S2A, Supplementary Table S5). Gene ontology analysis revealed 88 biological processes, which were involved in baseline refractive eye development (Supplementary Figure S2B, Supplementary Table S8). The gene ontology data highlighted an important role of visual perception, synaptic transmission, cell-cell communication, retina development, and DNA methylation in baseline refractive eye development. Moreover, baseline refractive development was associated with several canonical signaling pathways, including β-adrenergic signaling, protein kinase A signaling, dopamine receptor signaling, HIPPO signaling, mTOR signaling, phototransduction pathway, axonal guidance signaling, synaptic long-term potentiation, tight junction signaling, and DNA methylation signaling (Supplementary Figure S2C, Supplementary Table S11). We found that approximately 25% of genes involved in the regulation of visual acuity (268 out of 1,073 genes) were also associated with baseline refractive eye development (OR = 3.5, P = 3.391 × 10^-78^) (Figure 3A, Supplementary Table S13). These genes were involved in at least 47 biological processes and 31 molecular functions. The top 30 biological processes shown in Figure 3B (Supplementary Table S15) included such processes as homophilic cell adhesion, metabolic process, lipid metabolism, protein autophosphorylation, response to ischemia, visual perception, and neuron projection morphogenesis, i.e., many of the key processes underlying visual acuity (Figure 2B). The majority of these processes were suppressed in animals with high visual acuity and activated in animals with low visual acuity (Figure 3B). In agreement with these observations, approximately 55% of the canonical signaling pathways involved in the regulation of visual acuity were also involved in baseline refractive eye development (Figure 3C). Most notable pathways included protein kinase A signaling, RhoGDI signaling, melatonin signaling, senescence pathway, Wnt/Ca+ pathway, nNOS signaling, β-adrenergic signaling, synaptic long-term potentiation, dopamine-DARPP32 feedback signaling, α-adrenergic signaling, and mTOR Signaling.

**Fig. 3.**
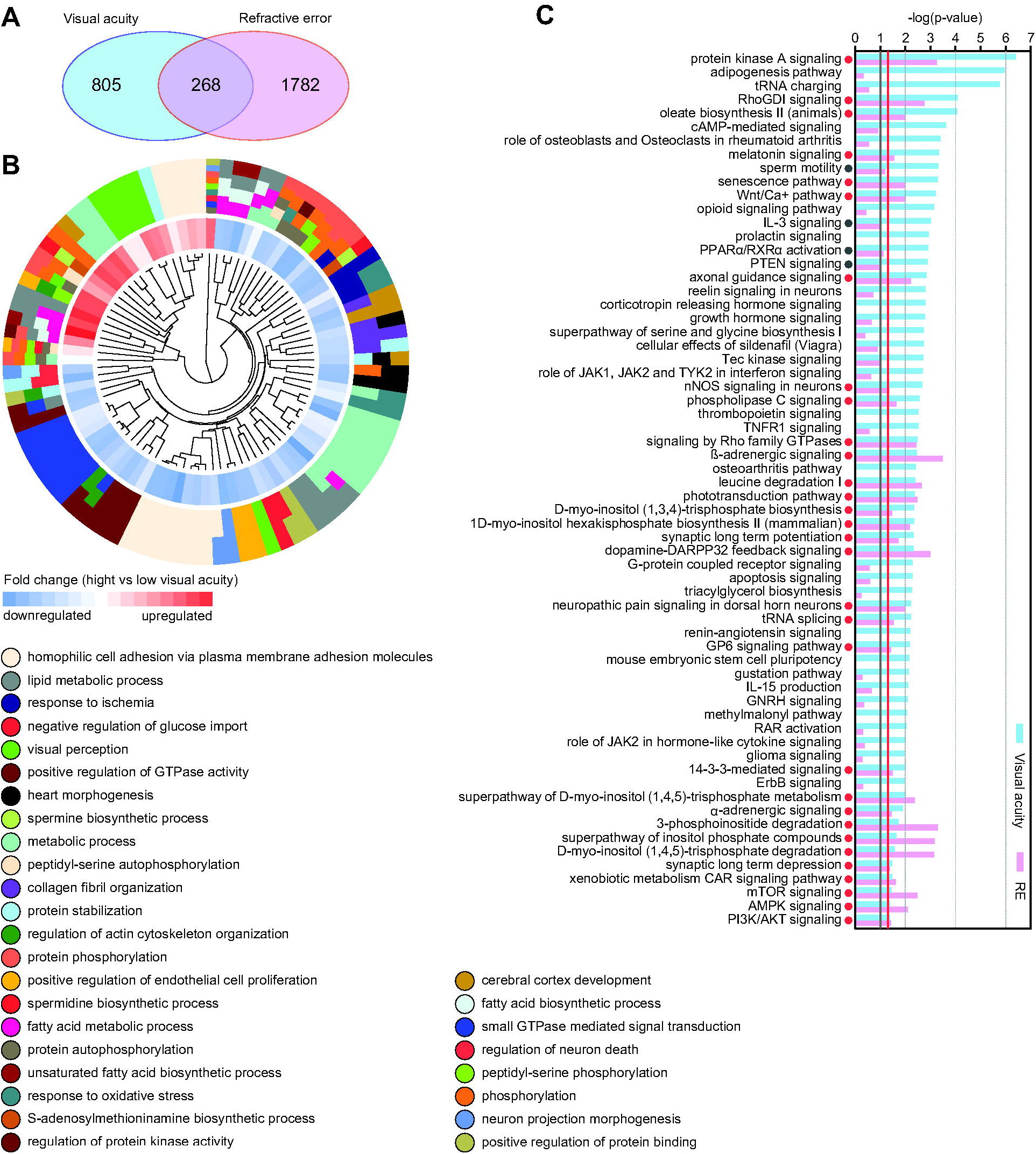
Gene expression network regulating visual acuity makes significant contribution to baseline refractive eye development. (A) Venn diagram showing overlap between genes regulating visual acuity and genes underlying baseline refractive eye development. (B) Top 30 biological processes affected by the genes involved in the regulation of visual acuity and baseline refractive development. Outer circle of the hierarchical clustering diagram shows hierarchical clusters of biological processes (identified by different colors); inner circle shows clusters of the associated genes up- or down-regulated in mice with high visual acuity versus mice with low visual acuity. (C) Canonical signaling pathways involved in the regulation of visual acuity and baseline refractive eye development. Vertical red line and red dots indicate P = 0.05. Vertical grey line and grey dots indicate P = 0.1. Colors identify pathways associated with either visual acuity or baseline refractive development and correspond to the colors in the Venn diagram (A).

Thus, although linear regression analysis demonstrated relatively modest influence of visual acuity on baseline refractive eye development, we found a substantial overlap between gene expression networks and canonical signaling pathways underlying these two processes.

### 3.4. Comparison of the transcriptome for visual acuity with the transcriptome for form-deprivation myopia reveals modest contribution of the signaling cascade subserving visual acuity to visually guided eye emmetropization

To examine the contribution of the genetic network regulating visual acuity to visually guided eye emmetropization, we analyzed the overlap between gene expression networks underlying visual acuity and susceptibility to form-deprivation myopia. As we reported previously [30], expression of 1,347 genes (Q-value < 0.01) was associated with susceptibility to form-deprivation myopia (Supplementary Figure S3A, Supplementary Table S6). Gene ontology analysis revealed involvement of several biological processes, including axonogenesis, transport, fatty acid oxidation, aging, insulin secretion, cell proliferation, cell-cell adhesion, and cellular response to hypoxia (Supplementary Figure S3B, Supplementary Table S9). Furthermore, susceptibility to form-deprivation myopia was linked to several canonical signaling pathways, including G2/M DNA damage checkpoint regulation, iron homeostasis signaling pathway, tight junction signaling, sirtuin signaling pathway, β-adrenergic and α-adrenergic signaling, HIPPO signaling, mTOR signaling, amyotrophic lateral sclerosis signaling, axonal guidance signaling, and growth hormone signaling (Supplementary Figure S3C, Supplementary Table S12). We discovered that approximately 19% of genes (205 out of 1,073 genes) involved in the regulation of visual acuity were also involved in the regulation of susceptibility to form-deprivation myopia (OR = 4.1, P = 5.219 × 10^-70^) (Figure 4A, Supplementary Table S14). These genes were involved in approximately 31 biological process and 19 molecular functions. As shown in Figure 4B (Supplementary Table S16), the top 30 biological processes included brain development, response to hydrogen peroxide, spermine and spermidine biosynthesis, metabolic process, response to ischemia, homophilic cell adhesion, regulation of autophagy, chloride transmembrane transport, and synaptic vesicle recycling. The absolute majority of these processes were suppressed in animals with high visual acuity and activated in animals with low visual acuity. Only 28% of canonical signaling pathways involved in the regulation of visual acuity were also involved in the regulation of susceptibility to form-deprivation myopia (Figure 4C). In addition to such pathways as tRNA charging, growth hormone signaling, methylmalonyl pathway, RAR activation pathway, role of JAK2 in hormone-like cytokine signaling, 2-oxobutanoate degradation, and iron homeostasis signaling pathway, these pathways included many of the same pathways involved in baseline refractive eye development, i.e., axonal guidance signaling, β-adrenergic signaling, GP6 signaling pathway, superpathway of D-myo-inositol (1,4,5)-trisphosphate metabolism, α-adrenergic signaling, 3-phosphoinositide degradation, superpathway of inositol phosphate compounds, D-myo-inositol (1,4,5)-trisphosphate degradation, mTOR signaling, and AMPK signaling.

**Fig. 4.**
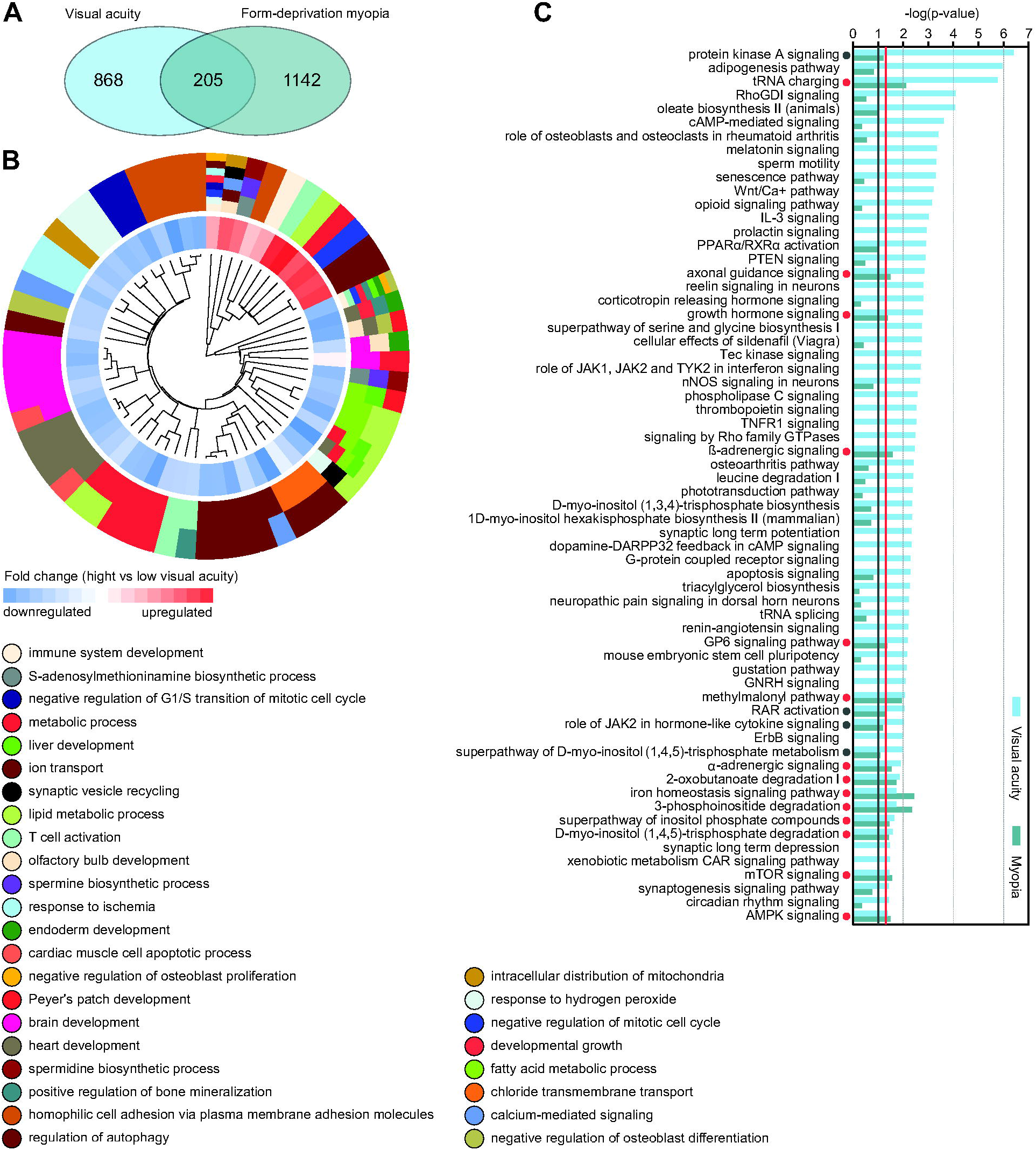
Gene expression network regulating visual acuity makes modest contribution to visually guided eye emmetropization. (A) Venn diagram showing overlap between genes regulating visual acuity and genes underlying form-deprivation myopia. (B) Top 30 biological processes affected by the genes involved in the regulation of visual acuity and susceptibility to form-deprivation myopia. Outer circle of the hierarchical clustering diagram shows hierarchical clusters of biological processes (identified by different colors); inner circle shows clusters of the associated genes up- or down-regulated in mice with high visual acuity versus mice with low visual acuity. (C) Canonical signaling pathways involved in the regulation of visual acuity and susceptibility to form-deprivation myopia. Vertical red line and red dots indicate P = 0.05. Vertical grey line and grey dots indicate P = 0.1. Colors identify pathways associated with either visual acuity or susceptibility to form-deprivation myopia and correspond to the colors in the Venn diagram (A).

In summary, these data suggest that the gene expression network regulating visual acuity has relatively limited contribution to visually guided eye emmetropization. Moreover, many of the canonical signaling pathways regulating both visual acuity and eye emmetropization are also involved in baseline refractive eye development.

### 3.5. Genes regulating visual acuity are involved in human myopia development and other eye-relatedgenetic disorders

Our data suggested that genes involved the regulation of visual acuity contributed to both baseline refractive eye development and visually guided eye emmetropization. Therefore, we than evaluated the association of these genes with human myopia revealed by GWAS and gene-based GWAS studies and other human eye-related genetic disorders listed in the Online Mendelian Inheritance in Man (OMIM) database. We found that 61 out of 381 genes involved in the regulation of visual acuity and refractive eye development was associated with known eye-related human disorders (Figure 5, Supplementary Table S17). Approximately 51% of these genes (31 gene) were associated with baseline refractive eye development, 33% (20 genes) were involved in visually guided emmetropization, and 16% (10 genes) were associated with both baseline refractive eye development and emmetropization. The largest number of genes implicated in human disorders was linked to myopia (57%), whereas other genes were associated with retinal degeneration and retinal atrophy, neurodevelopmental disorders, and connective tissue disorders. The most prevalent biological processes affected by these genes were visual perception, calcium ion transport, metabolic processes, and neurogenesis (Supplementary Table S18).

**Fig. 5.**
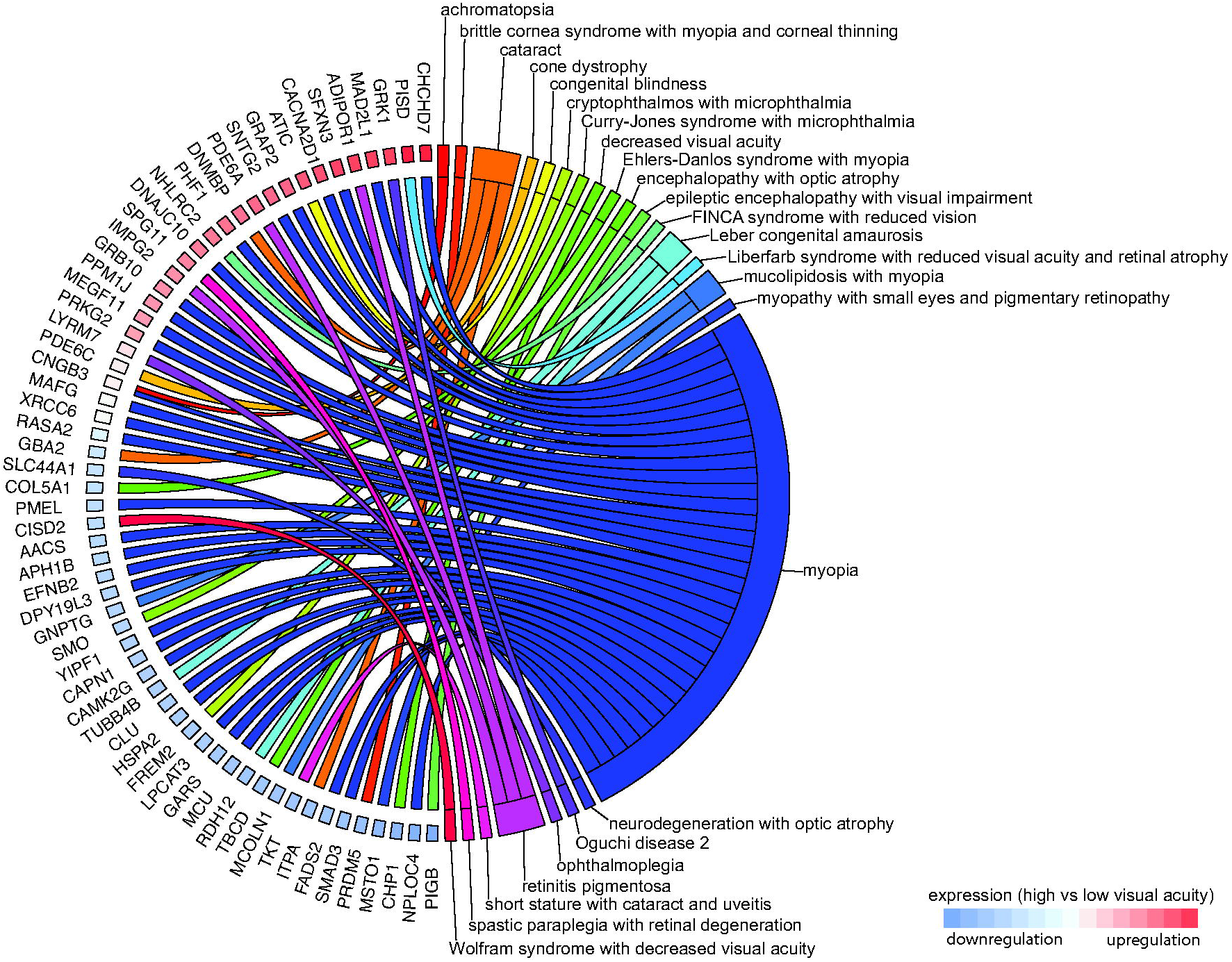
Genes involved in the regulation of visual acuity are associated with the development of myopia and other human genetic disorders. Chord diagram showing genes (left semicircle) and human genetic disorders (right semicircle) associated with visual acuity regulation and refractive eye development in mice. Colored bars underneath gene names show up- or down-regulation of the corresponding genes in mice with high visual acuity versus mice with low visual acuity.

Cumulatively, these data suggest that the genetic network regulating visual acuity makes a substantial contribution to the development of human myopia by modulating a number of biological processes involved in refractive eye development, including phototransduction-related signaling, signal transduction, and signaling pathways underlying various metabolic processes.

## 4. Discussion

Our recent data suggest that baseline refractive eye development and visually guided eye emmetropization are controlled by overlapping but largely distinct genetic networks [28]. We also found that the genetic network subserving contrast perception has a significant contribution to optical defocus detection and emmetropization, while having a very little contribution to baseline refractive eye development. Growing evidence indicates that detection of optical defocus by the retina is mediated by luminance contrast and longitudinal chromatic aberrations [8, 86]; however, some findings suggest that signaling cascades regulating visual acuity may also be involved in eye emmetropization [67]. Our current data suggest that the retinal gene expression network subserving visual acuity has a limited but significant involvement in both baseline refractive eye development and visually guided eye emmetropization. However, contribution of the genetic network underlying visual acuity to baseline refractive eye development appears to be both different and more significant compared to its contribution to eye emmetropization.

We found that visual acuity is regulated by a large number of genes organized into a rather elaborate genetic network contributing to a variety of biological processes, molecular functions and signaling pathways. As expected, our data clearly suggest that visual acuity depends on the density of photoreceptors because biological processes related to nervous system development and positive regulation of cell proliferation were significantly overrepresented. However, visual acuity seems to be also associated with transcriptional regulation of gene expression, protein phosphorylation, and protein turnover, suggesting that visual acuity is also actively modulated via dynamic changes in gene expression and protein signaling networks, beyond a sheer number of photoreceptors per unit of retinal surface. Overrepresentation of biological processes related to cell adhesion and axonogenesis points to a prominent role of synaptic transmission in the regulation of visual acuity. We found that high visual acuity was associated with activation of the RhoGDI signaling and PPARα/RXRα activation pathways and strong suppression of the protein kinase A signaling pathway, oleate biosynthesis, opioid signaling pathway, reelin signaling in neurons, Tec kinase signaling, and β-adrenergic signaling. Although the majority of these biological processes and signaling pathways were implicated in the regulation of visual acuity for the first time, the opioid signaling pathway was previously shown to modulate visual acuity [28]. For example, cannabinoids were shown to influence visual acuity through the cannabinoid receptor type 2 (CB2R), which closely interacts with the opioid signaling pathway [77, 87–90]. Noteworthy, expression of *CB2R* was shown to be regulated by nitric oxide, which has long been implicated in refractive eye development [12, 91–95].

Although our linear regression analyses suggested that visual acuity has essentially equal contribution to both baseline refractive eye development and visually guided eye emmetropization, analysis of gene expression networks revealed a more complex relationship between visual acuity, baseline refractive development, and emmetropization. Although 92 out of 268 genes involved in the regulation of both visual acuity and baseline refractive development were also implicated in emmetropization (OR = 48.2; P = 1.416 × 10^-135^), we found that the key biological processes and signaling pathways impacted by the visual-acuity-related genetic network in baseline refractive development and visually guided eye emmetropization were substantially different. While visual-acuity-related biological processes involved in baseline refractive development were primarily related to cell-cell communication, metabolic processes, GTPase-mediated signaling, visual perception, and neuron projection morphogenesis, visual-acuity-related biological processes involved in emmetropization were primarily related to neurogenesis, ion transport, metabolic processes, cell-cell communication, response to ischemia, and response to hydrogen peroxide. The visual-acuity-related genetic network contributes to several canonical pathways common to both baseline refractive development and emmetropization, including pathways involved in inositol metabolism, protein kinase A signaling, α-adrenergic and β-adrenergic signaling, axon guidance, GP6 signaling, mTOR signaling, and AMPK signaling. However, we also uncovered substantial differences. First, the total number of canonical pathways underlying baseline refractive development and encoded by the visual-acuity-related genetic network is much greater compared to the number of pathways underlying eye emmetropization and encoded by the visual-acuity-related genetic network (35 versus 18). Second, despite the existence of shared pathways, the visual-acuity-related genetic network contributes to largely different canonical pathways when it comes to baseline refractive development and visually guided eye emmetropization. Most notable pathways unique to baseline refractive development include melatonin signaling, senescence pathway, Wnt/Ca+ pathway, nNOS signaling, phototransduction pathway, synaptic long-term potentiation, synaptic long-term depression, and dopamine-DARPP32 feedback signaling. Conversely, pathways unique to visually guided eye emmetropization are limited to tRNA charging, growth hormone signaling, methylmalonyl pathway, RAR activation, JAK2 signaling, 2-oxobutanoate degradation, and iron homeostasis signaling pathway. Taken together, these data suggest that the visual-acuity-related gene network makes largely unique contributions to the retinal signaling cascades underlying baseline refractive eye development and visually guided eye emmetropization.

Several biological processes and signaling pathways regulating visual acuity, baseline refractive eye development and eye emmetropization were previously implicated in refractive eye development. For example, cell-cell communication and synaptic transmission were shown to influence refractive eye development and response to optical defocus by several studies [12, 28, 29, 96–99]. Energy production and various metabolic processes were also shown to play an important role in refractive eye development and eye’s response to optical defocus [12, 28, 29, 98–100]. The protein kinase A signaling, α-adrenergic signaling, β-adrenergic signaling, and mTOR signaling pathways have been repeatedly found to be linked to retinal signaling underlying refractive eye development [12, 28, 98, 100].

The biological processes and signaling pathways uniquely underlying either baseline refractive development or visually guided eye emmetropization and regulated by the visual-acuity-related genetic network provide several interesting insights into the biology of refractive eye development. The finding that the genetic network regulating visual acuity and baseline refractive development contributes to melatonin signaling, senescence pathway, phototransduction pathway, and synaptic long-term depression reveals a complex interplay between genetic networks underlying visual acuity, perception of contrast, and baseline refractive development because it was found that genes controlling both contrast perception and baseline refractive development encode components of the same pathways. Interestingly, the tRNA charging and JAK2 signaling pathways involved in both visual acuity regulation and visually guided eye emmetropization are also involved in contrast perception, thus suggesting an overlap between genetic networks underlying visual acuity regulation, contrast perception, and visually guided eye emmetropization.

We discovered that approximately 16% of the genes, which we found to be involved in visual acuity regulation and refractive eye development, were also implicated in human myopia and other eye-related genetic disorders. Several genes were involved in visual perception. One of the genes, *GRK1,* causing Oguchi disease is involved in light-dependent deactivation of rhodopsin [101]. The myopia-associated gene *CNGB3* encodes one of the subunits that form cyclic nucleotide-gated ion channels required for sensory transduction in rod photoreceptors [28, 102]. The *IMPG2* gene linked to retinitis pigmentosa plays a role in organization of the interphotoreceptor matrix and promotes growth and maintenance of the photoreceptor outer segment [103, 104]. The two genes, *PDE6C* known to cause cone dystrophy and *PDE6A* linked to retinitis pigmentosa, encode subunits of cone and rod phosphodiesterases which play important roles in phototransduction [105–109]. The *RDH12* gene involved in the development of Leber congenital amaurosis encodes retinol dehydrogenase which metabolizes both all-trans- and cis-retinols, which play a key role in the phototransduction pathway [110, 111]. The two myopia-associated genes *CACNA2D1* and *CAMK2G* encode several subunits of the volta-gegated calcium channel and a calcium-dependent protein kinase, which play important roles in the regulation of synaptic transmission in the retina downstream of photoreceptors [28, 112–115]. Interestingly, calcium signaling in the retina was shown to be modulated by dopamine and nitric oxide, both of which have been implicated in refractive eye development [92–95, 116–122]. Taken together, these data suggest that visual acuity depends, to a significant degree, on the signaling cascade underlying light perception and phototransduction. While deleterious mutations in these genes often cause severe retinal damage associated with photoreceptor degeneration, genetic variations modulating gene expression appear to primarily influence visual acuity and refractive eye development.

## 5. Conclusions

In this study, we explored the genetic network regulating visual acuity and its contribution to baseline refractive eye development and visually guided emmetropization. Our results provide evidence that visual acuity is determined not only by the density of photoreceptors per unit of retinal surface, but it is also regulated at the level of molecular signaling in the retina by an extensive genetic network. This genetic network controls visual acuity via a diverse set of biological processes primarily related to transcriptional regulation of gene expression, protein phosphorylation, protein turnover, homophilic cell adhesion, metabolic processes, axonogenesis, response to ischemia, photoreceptor signaling, and synaptic transmission. Our data suggest that the genetic network regulating visual acuity makes modest but significant contributions to both baseline refractive eye development and visually guided eye emmetropization; however, the contribution to baseline refractive eye development is much more substantial. We found that visual-acuity-related pathways uniquely involved in baseline refractive eye development include melatonin signaling, senescence pathway, Wnt/Ca+ pathway, nNOS signaling, phototransduction pathway, synaptic long-term potentiation, synaptic long-term depression, and dopamine-DARPP32 feedback signaling. Visual-acuity-related pathways uniquely underlying visually guided eye emmetropization are primarily involved in tRNA charging, growth hormone signaling, methylmalonyl pathway, RAR activation, JAK2 signaling, 2-oxobutanoate degradation, and iron homeostasis signaling. A small number of genes involved in the regulation of visual acuity are also involved in the development of human myopia. These genes are overwhelmingly involved in light perception and phototransduction. Our results also suggest a complex interplay between genetic networks that control visual acuity, contrast perception, and refractive eye development.

## Availability of data and materials

All data generated or analyzed during this study are included in this article and its supplementary information files. The RNA-seq data have been deposited in the Gene Expression Omnibus database [GSE158732] (https://www.ncbi.nlm.nih.gov/geo/query/acc.cgi?acc=GSE158732).

## Author contributions

TVT and AVT conceptualized the study, analyzed refractive eye development, susceptibility to form-deprivation myopia and visual acuity in mice, performed RNA-seq, and analyzed the data. AVT supervised the entire study, analyzed and validated data, and wrote the original draft of the manuscript. All authors read, edited, and approved the final version of the manuscript.

## Abbreviations

AMPK: AMP-activated protein kinase
ANOVA: analysis of variance
ARVO: Association for Research in Vision and Ophthalmology
CACNA2D1: calcium voltage-gated channel auxiliary subunit alpha2 delta 1
CAMK2G: calcium/calmodulin dependent protein kinase II gamma
CB2R: cannabinoid receptor type 2
CFH: complement factor H
CNGB3: cyclic nucleotide gated channel subunit beta 3
cpd: cycles per degree
DARPP32: protein phosphatase 1 regulatory inhibitor subunit 1B
DAVID: database for annotation, visualization and integrated discovery
GP6: glycoprotein VI platelet
GRK1: G protein-coupled receptor kinase 1
GWAS: genome-wide association study
IMPG2: interphotoreceptor matrix proteoglycan 2
IPA: Ingenuity Pathway Analysis
JAK2: Janus kinase 2
LED: light-emitting diode
LRIT1: leucine rich repeat, Ig-like and transmembrane domains 1
mTOR: mammalian target of rapamycin
nNOS: neuronal nitric oxide synthase
OMIM: Online Mendelian Inheritance in Man database
OR: odds ratio
PDE6A: phosphodiesterase 6A
PDE6C: phosphodiesterase 6C
PPARα: Peroxisome proliferator-activated receptor alpha
PTEN: phosphatase and tensin homolog
RAR: retinoic acid receptor
RCB: randomized complete block
RDH12: retinol dehydrogenase 12
RhoGDI: Rho GDP-dissociation inhibitor
RIN: RNA integrity number
RNA-seq: massive parallel RNA sequencing
RXRα: retinoid X receptor alpha
SNP: single-nucleotide polymorphism
QTL: quantitative trait locus
VEP: visually evoked potential

## Supplementary materials

**Supplementary Tables: Table S1.** Visual acuity in different strains of mice (P40, cpd). **Table S2.** Baseline refractive errors in different strains of mice (P40, diopters). **Table S3.** Form-deprivation myopia in different strains of mice (deprived eye versus control eye, diopters). **Table S4.** Genes whose expression correlates with visual acuity in mice (FC = fold change, high versus low visual acuity). **Table S5.** Genes whose expression correlates with baseline refractive errors in mice (FC = fold change, hyperopia versus myopia). **Table S6.** Genes whose expression correlates with susceptibility to myopia in mice (FC = fold change, high versus low myopia). **Table S7.** Gene ontology categories associated with genes whose expression correlates with visual acuity in mice (BP, biological process; CC, cellular component; MF, molecular function). **Table S8.** Gene ontology categories associated with genes whose expression correlates with baseline refractive errors in mice (BP, biological process; CC, cellular component; MF, molecular function). **Table S9.** Gene ontology categories associated with genes whose expression correlates with susceptibility to myopia in mice (BP, biological process; CC, cellular component; MF, molecular function). **Table S10.** Canonical signaling pathways associated with genes whose expression correlates with visual acuity in mice. **Table S11.** Canonical signaling pathways associated with genes whose expression correlates with baseline refractive errors in mice. **Table S12.** Canonical signaling pathways associated with genes whose expression correlates with susceptibility to myopia in mice. **Table S13.** Genes whose expression correlates with both visual acuity and baseline refractive errors in mice (FC = fold change, visual acuity = high versus low visual acuity, RE = hyperopia versus myopia). **Table S14.** Genes whose expression correlates with both visual acuity and susceptibility to myopia in mice (FC = fold change, visual acuity = high versus low visual acuity, myopia = high versus low myopia). **Table S15.** Gene ontology categories associated with genes whose expression correlates with both visual acuity and baseline refractive errors in mice (BP, biological process; CC, cellular component; MF, molecular function). **Table S16.** Gene ontology categories associated with genes whose expression correlates with both visual acuity and susceptibility to myopia in mice (BP, biological process; CC, cellular component; MF, molecular function). **Table S17.** Visual acuity genes associated with refractive eye development in mice and linked to eye-related human diseases (FC = fold change, visual acuity = high versus low visual acuity, RE = expression correlates with RE, myopia = expression correlates with susceptibility to myopia). **Table S18.** Gene ontology categories associated with genes whose expression correlates with visual acuity, baseline refractive errors, and form-deprivation myopia and which are involved in human diseases (see Table S17) (BP, biological process).

**Supplementary Figure S1.** Baseline refractive error correlates weakly with susceptibility to form-deprivation myopia. Linear regression showing weak correlation between baseline refractive error and susceptibility to myopia. r, Pearson’s correlation coefficient; P, Pearson’s correlation significance.

**Supplementary Figure S2.** Gene expression network underlying baseline refractive eye development controls diverse biological functions and signaling pathways. (A) Expression of 2,050 genes (Q-value < 0.01) correlates with baseline refractive errors. Hierarchical clustering shows that differential genes are organized in two clusters: one (top) cluster exhibits increased expression in the myopic mice, and the second (bottom) cluster shows increased expression in the hyperopic mice. (B) Top 30 biological processes affected by the genes involved in baseline refractive eye development. Outer circle of the hierarchical clustering diagram shows hierarchical clusters of biological processes (identified by different colors); inner circle shows clusters of the associated genes up- or down-regulated in hyperopic mice versus myopic mice. (C) Top 30 canonical pathways affected by the genes associated with baseline refractive eye development. Horizontal yellow line indicates P = 0.05. Z-scores show activation or suppression of the corresponding pathways in animals with hyperopia versus animals with myopia.

**Supplementary Figure S3.** Gene expression network subserving susceptibility to form-deprivation myopia controls diverse biological functions and signaling pathways. (A) Expression of 1,347 genes (Q-value < 0.01) correlates with susceptibility to form-deprivation myopia. Hierarchical clustering shows that differential genes are organized in two clusters: one (top) cluster exhibits increased expression in mice with low susceptibility to myopia, and the second (bottom) cluster shows increased expression in mice with high susceptibility to myopia. (B) Top 30 biological processes affected by the genes involved in the development of form-deprivation myopia. Outer circle of the hierarchical clustering diagram shows hierarchical clusters of biological processes (identified by different colors); inner circle shows clusters of the associated genes up- or down-regulated in mice with high susceptibility to myopia versus mice with low susceptibility to myopia. (C) Top 30 canonical pathways affected by the genes associated with the development of form-deprivation myopia. Horizontal yellow line indicates P = 0.05. Z-scores show activation or suppression of the corresponding pathways in animals with high susceptibility to form-deprivation myopia versus animals with low susceptibility to form-deprivation myopia.

